# Deep Learning Characterization of Brain Tumours With Diffusion Weighted Imaging

**DOI:** 10.1101/2022.01.25.477747

**Authors:** Cameron Meaney, Sunit Das, Errol Colak, Mohammad Kohandel

**Affiliations:** Department of Applied Mathematics, University of Waterloo, Waterloo, Canada; Division of Neurosurgery, Li Ka Shing Knowledge Institute, St. Michael’s Hospital, Toronto, Canada; Faculty of Medicine, University of Toronto, Toronto, Canada; Department of Medical Imaging and Li Ka Shing Knowledge Institute, St. Michael’s Hospital, Toronto, Canada; Odette Professorship in Artificial Intelligence for Medical Imaging, St. Michael’s Hospital, Toronto, Canada

## Abstract

Glioblastoma multiforme (GBM) is one of the most deadly forms of cancer. Methods of characterizing these tumours are valuable for improving predictions of their progression and response to treatment. A mathematical model called the proliferation-invasion (PI) model has been used extensively in the literature to model these tumours, though it relies on known values of two key parameters: the tumour cell diffusivity and proliferation rate. Unfortunately, these parameters are difficult to estimate in a patient-specific manner, making personalized tumour projections challenging. In this paper, we develop and apply a deep learning model capable of making accurate estimates of these key GBM-characterizing parameters while simultaneously producing a full projection of the tumour progression curve. Our method uses two sets of multi sequence MRI imaging in order to make predictions and relies on a preprocessing pipeline which includes brain tumour segmentation and conversion to tumour cellularity. We apply our deep learning model to both synthetic tumours and a dataset consisting of five patients diagnosed with GBM. For all patients, we derive evidence-based estimates for each of the PI model parameters and predictions for the future progression of the tumour. Discussion and implications for future work and clinical relevance are included.

## 1 Introduction

The most prevalent form of malignant brain tumour among adults is glioblastoma multiforme (GBM). Modern treatments for GBMs involve a combination of surgery, chemotherapy, and radiotherapy, yet despite the most aggressive therapies, mean survival after diagnosis is only slightly over a year [3, 36]. GBMs are commonly distinguished from other brain tumours by their dense necrotic core and a surrounding shell of peritumoral edema, both of which pose challenges for effective therapy. In spite of continuous advancements in our understanding of the biology of GBMs and in the imaging techniques used to observe them, their full extent is often inaccurately assessed by state of the art imaging: many studies have noted that their growth and invasion tends to be underestimated. As treatments are typically designed based on this knowledge, there is potential for therapies to be misapplied, possibly contributing to recurrence or treatment resistance. Methods and technologies for more accurately assessing GBM growth and invasion are sought by clinicians and researchers, with the hope of using this knowledge to make more accurate growth projections and design more effective treatments. Furthermore, if improved methods allowed for recommendations to be made in a patient specific manner, then personalized medicine for GBMs becomes closer to reality.

Much of the information available when examining GBMs comes from medical imaging, specifically multi sequence magnetic resonance imaging (MRI). Qualitative conclusions can be drawn from MRIs following analysis which can be used by clinicians to inform recommendations for observation and treatment. But with the explosion of big data in recent years, many have begun to analyze medical imaging data quantitatively, which has proven tremendously fruitful. Unfortunately, the barrier to entry in quantitative analysis of medical imaging is quite high, as it requires the use of substantial mathematical and computational tools in order to draw relevant and actionable conclusions. Understandably, mathematical modelling and machine learning have become crucial tools with the capability to make quantitative analysis of imaging data accessible enough for use in a clinical setting.

Many works have used mathematical models to predict the extent, future growth, and response to treatment of GBMs [1, 11,13, 21, 22, 25, 26, 41]. In particular, the most common model used to simulate GBM progression is a partial differential equation (PDE) called the proliferation-invasion (PI) model. The PI model is a relatively simple model which describes GBM progression being governed by two distinct processes: diffusion and proliferation. Each of these processes is characterized by a model parameter that dictates the importance of that process to the overall progression. With estimates of the values of these two parameters, the PI model is complete and can be used to make projections of tumour progression and response to treatment. However, making accurate estimates of these two parameters is quite challenging. Given the amount of data that standard numerical tools require, the costs, difficulties, and potential risks associated with obtaining the necessary imaging data are often disqualifying. Researchers must then resort to techniques capable of making predictions based on smaller data sets which, while possible, often introduces a new set of challenges and inaccuracies.

A method does exist for estimating these parameters, though it comes with its own strengths and weaknesses. Swanson et al. described this technique in a paper [40] and subsequent patent [39] and it has remained the dominant method since its inception. While beautiful in its simplicity and ingenuity, it relies on a few ideas which dull its accuracy; specifically, potentially unreasonable mathematical assumptions, sensitivity to input noise, and reliance on human measurement. Additionally, this method fails to make use of all of the data available to it, opting to take only a few measurements from the orders of magnitude more that exist. A method which eliminates these weaknesses and utilizes more of the available data could provide more precise estimates than were otherwise possible, allowing for more accurate projections of progression and treatment response.

In this paper, we develop a specialized deep learning model capable of characterizing brain tumours by making more accurate estimations of the PI model parameters. Our method requires multi sequence MRI including the T1, T1-GAD, T2, T2-FLAIR, and DWI sequences at two time points without treatment between and produces estimates of the PI model parameters along with a full projection of the tumour progression over time. Involved in our pipeline are several important steps including segmentation, conversion to tumour cellularity, and network training; each of which are described in the methods section below. In the results section, we first exhibit the capabilities of our technique by applying it to ‘synthetic’ tumours for which we know the ground truth and can evaluate the accuracy. We then apply our method to real patient GBM imaging data and derive patient-specific estimates and growth projections. In the conclusion section, the work is summarized, limitations are discussed, and directions for future research are noted.

## 2 Methods

This paper outlines a deep learning model capable of producing accurate, patient-specific estimates of the PI model parameters and simultaneously giving a full projection of the GBM progression curve. As input, the neural network requires the voxel-by-voxel tumour cellularity throughout the tumour volume at each imaging time. The following sections detail the key steps and ideas involved in deriving these cellularity profiles from MRI data, including data acquisition, preprocessing, segmentation, and conversion. We also explain and discuss the PI model and thoroughly describe the deep learning model used to make predictions. Our procedure pipeline can also be seen in figure 1.

**Figure 1:**
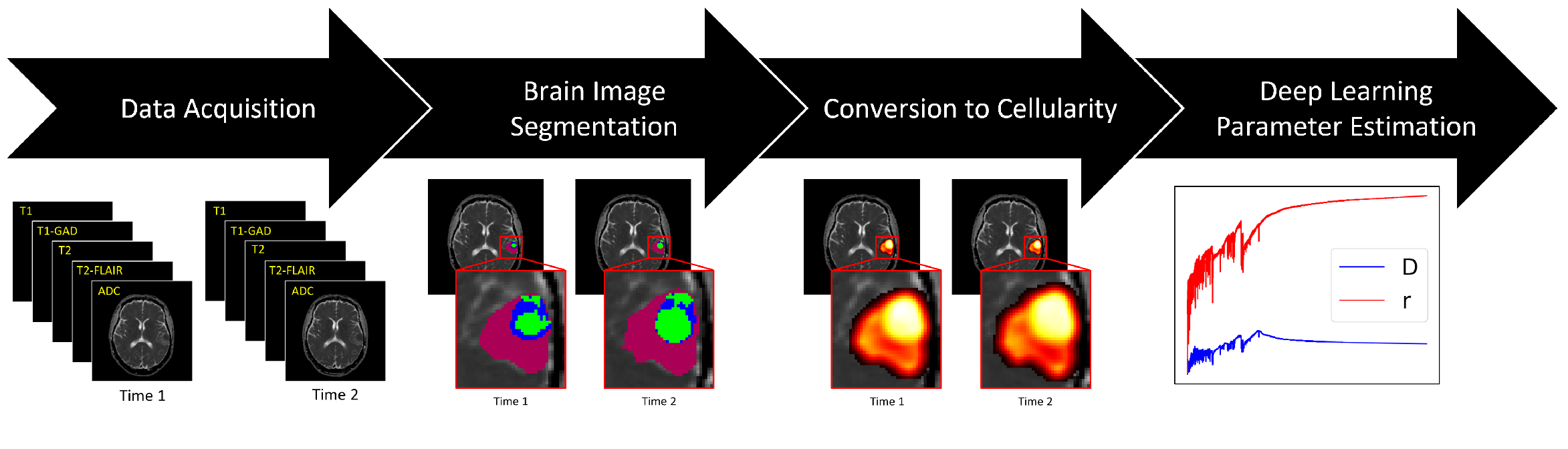
Pipeline for our parameter estimation model. Following data acquisition, the T1, T1-GAD, T2, and T2-FLAIR images are used for segmenting the tumour voxel-by-voxel. The segmentation results are used to convert the ADC to a map of tumour cellularity. The resulting tumour cellularities at the two imaging times are given as input to our deep learning model to derive PI model parameter estimates and progression curve projections.

### 2.1 Proliferation Invasion Model

The mathematical model most commonly used to describe the progression of GBMs is a PDE called the proliferation-invasion model which has been used and discussed in previous works [1, 11, 13, 21, 22,25, 26, 41]. The PI model has the form

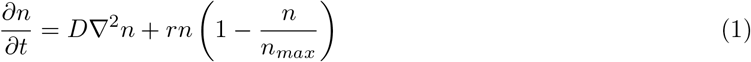

where 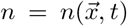 represents the tumour cell density at position 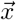 and time *t*. The key characterizing parameters of the PI model are *D* and *r*, which represent the tumour cell diffusivity and proliferation rate respectively. With estimates of these two parameters, the PI model can be solved and the tumour fully characterized. Also present in the model is the cell carrying capacity *n*_*max*_, which quantifies the maximum biologically feasible tumour cellularity at a location. Typically, the value of *n*_*max*_ is straightforward to estimate as is simply assumed to be the maximum observed cell density. Furthermore, when using the PI model in our model, we work with a nondimensionalized form of the equation which considers only the ratio 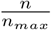. For this reason, we artificially set *n*_*max*_ = 1 for all calculations without loss of generality. In our nondimensionalization, we scale the time variable by the time between images and the spatial variable by the image voxel size.

### 2.2 Data Acquisition

The imaging data used in our study was collected as part of previous studies and was obtained for use in this work through a data sharing agreement between the University of Waterloo and Unity Health Toronto. Research ethics board approval was obtained from both the University of Waterloo and Unity Health Toronto and all ethics guidelines were followed in the study. The dataset consists of MRI data stored in DICOM format for five patients, each of whom was diagnosed with GBM. For each patient, MRI was performed in two instances with the T1, T1-GAD, T2, T2-FLAIR, and DWI sequences collected, giving 10 sequences per patient, each of which is 3D and voxelized. As explained below, the first four of these sequences are used in a segmentation algorithm. The DWI, however, is used to derive a map of the tumour cellularity. Importantly, what is actually required for this derivation of cellularity is the map of apparent diffusion coefficient (ADC) which is often obtained explicitly but can also be derived from a DWI and the T2 (null, *b* = 0) image. A summary of our dataset can be seen in table 1.

**Table 1:** Summary of the patient data used in this study.

| Patient ID | Days Between Imaging | Diagnosis |
| --- | --- | --- |
| 001 | 121 | GBM grade IV |
| 002 | 26 | GBM grade IV |
| 003 | 51 | GBM grade IV |
| 004 | 111 | GBM grade IV |
| 005 | 76 | Anaplastic astrocytoma, progressed to GBM grade IV |

As our study aims to characterize the natural progression of the tumour, a requirement for our data was that no anticancer treatment was performed between the imaging times for each patient. The only required data for our study are the images and time between them; therefore, the dataset is de-identified with respect to all patient identifiers.

### 2.3 Segmentation

The next step in our model is to segment the tumour from the brain images. Brain tumour segmentation from imaging is a well-studied problem in the literature - even a cursory search will return myriad results with countless segmentation approaches to consider. In this work, we rely on the Federated Tumor Segmentation (FeTS) initiative software which is an open source toolkit which utilizes machine learning techniques to perform tumour boundary detection [27, 34, 35] (https://www.fets.ai/). It is developed and maintained by the Centre for Biomedical Image Computing and Analytics at the University of Pennsylvania. The software utilizes a fusion of numerous deep learning models to complete its segmentation. More specifically, the FeTS segmentation algorithm uses T1, T1-GAD, T2, and T2-FLAIR 3D sequences and classifies each voxel into one of four tissue categories: peritumoral edema, enhancing proliferative, necrotic, or non-tumoral. As is explained below, this classification into four tissue types rather than the simple binary classification of tumoral vs. non-tumoral enables a more sophisticated conversion to cellularity.

Our original DICOM images were first converted to NIfTI files as required by FeTS. Then for each time and patient respectively, the T1, T2, T2-FLAIR, and DWI images were co-registered to the T1-GAD image. As part of this co-registration process, the 3D voxel size on all images was standardized to (1mm, 1mm, 1mm). Representative cross sections of the resulting segmentations can be seen in figure 2.

**Figure 2:**
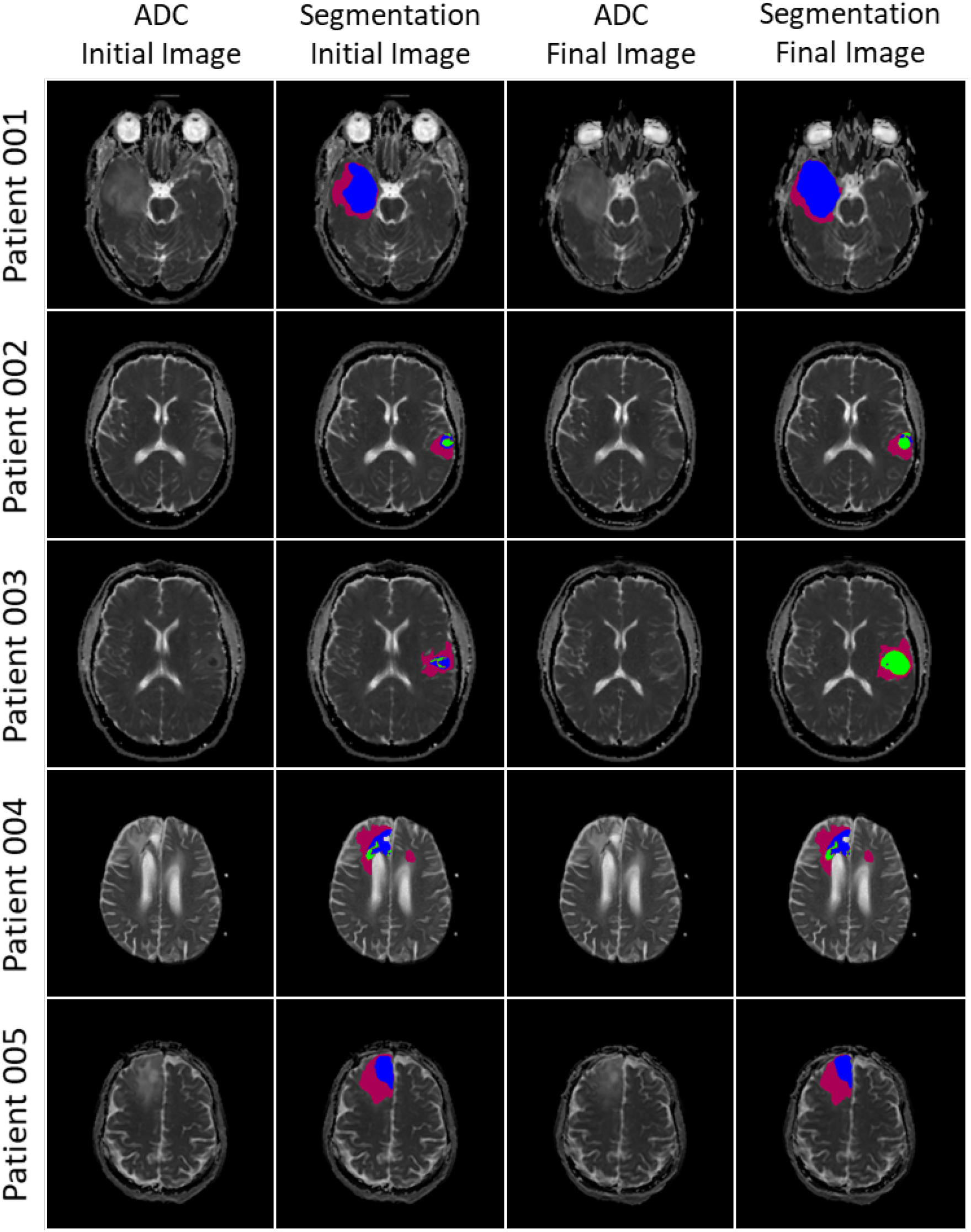
Representative cross sections of segmentation results for our patient dataset. Segmentaions were performed by the FeTS software which uses the T1, T1-GAD, T2, and T2-FLAIR images to classify the image voxel-by-voxel into the tissue categories peritumoral edema, enhancing proliferative, necrotic, or nontumoral. The first and third columns show the ADC at the initial and final image times respectively. The second and fourth columns show the ADC with the segmentation results superimposed at the initial and final image times respectively. Green voxels represent necrotic areas, blue voxels enhancing proliferative areas, and pink voxels peritumoral edema areas.

### 2.4 Conversion to Tumour Cell Density

The input required for our deep learning model is the tumour cellularity at each imaging time. Plenty of literature exists discussing methods to obtain a profile of tumour cell density from imaging, the most popular of which relies upon the ADC image. Many works have examined the correlation between ADC and tumour cell density, specifically noting that

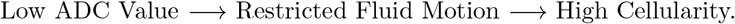

The biological logic behind this line of thinking is that the largest barrier to the Brownian motion of water molecules present in the tissue is cell membranes, and that an increase in cellularity prevents the free diffusion of water molecules observed on an ADC image. Furthermore, this thinking is supported by more than simply a biological rationale, with many experimental works reporting strong correlation between ADC and tumour cellularity, particularly in gliomas [8, 10, 12, 16, 37, 43, 45]. Most convincingly, Surov et al. [38] provided a meta-analysis of the correlations between ADC and cellularity for various types of tumours, noting that gliomas exhibited one of the highest correlations over many cancer types. This correlation has led to the development of a mathematical expression relating the local ADC and tumour cellularity, which is given by

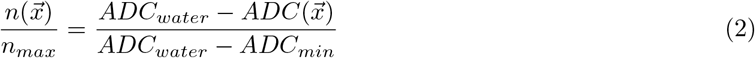

where *ADC*_*water*_ is the ADC in free water, *ADC*_*min*_ is the minimum ADC observed in the tumour, and *n*_*max*_ is the tumour cell carrying capacity also found in the PI model (equation (1)). This expression assumes a simple negative linear relationship between ADC and cellularity with the maximum cellularity coinciding with the minimum observed ADC and the minimum cellularity coinciding with the maximum observed ADC, that of water. Many previous mathematical works have used this relationship to derive tumour cell density profiles [4, 5, 23, 42].

While this relationship has the benefits of being simple and experimentally validated, it can occasionally lead to cell density profiles that are not biologically reasonable. For example, in a recent review of diffusion imaging of brain tumours, Maier et al. [17] noted that ADC values in brain tumour tissue clearly exceed ADC values of normal gray and white matter, and furthermore that ADC values tend to be highest in the centre of the tumour and decrease toward the tumour boundary. Using this mathematical conversion relationship for glioblastoma would therefore result in a cellularity profile highest at the tumour boundary and lowest at the tumour core. Of course, this is the opposite of the biology expected of GBMs which, particularly in cases of central necrosis, are typically known for their highly cellular core. Maier et al explain this apparent contradiction by explaining that the increased ADC observed in the tumour core is the result of the degradation of cell membranes common in necrotic tissues [17]. Another potential issue is that due to the sharp nature of tumour segmentation maps (tumour or not tumour), the cell density profile resulting from this conversion will have a sharp drop-off at the tumour boundary. Again, this is not the biology of GBMs, which tend to invade local tissues with a smaller number of cells than is present in the tumour core. Furthermore, both of these drawbacks lead to results that are opposite of the results typical of the PI model, which produces profiles with a dense core and decreasing cellularity toward the tumour boundary.

However, the additional knowledge obtained from using FeTS for segmentation allows for the creation of a more sophisticated conversion relationship. Specifically, consider a voxelwise conversion function as follows.

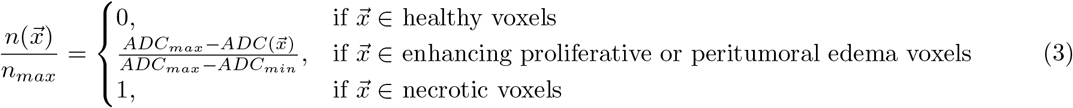

In this relationship, the additional knowledge from classes of tissue identified by FeTS is leveraged to ensure that the resulting cellularity profiles agree with GBM biology and the PI model. It is assumed that in necrotic tissue, the cells have ceased proliferating and have therefore reached their carrying capacity, *n*_*max*_. In proliferative or edematous voxels, the established relationship is used, and in healthy tissues the cellularity is assumed to be zero. While this form alleviates the problem of low cellularity in the tumour core, it does not address the sharp discontinuities occurring at the boundary between tissue types. To address this, a mean filter is applied after conversion to smooth the boundaries. Though this new conversion model is not perfect, it is able to remove many of the problems with the existing conversion by incorporating information from a more sophisticated segmentation algorithm.

In our derivation of cellularity from multi sequence MRI, we use this conversion procedure. The cell densities resulting from this conversion method for our patient data can be seen in figure 3.

**Figure 3:**
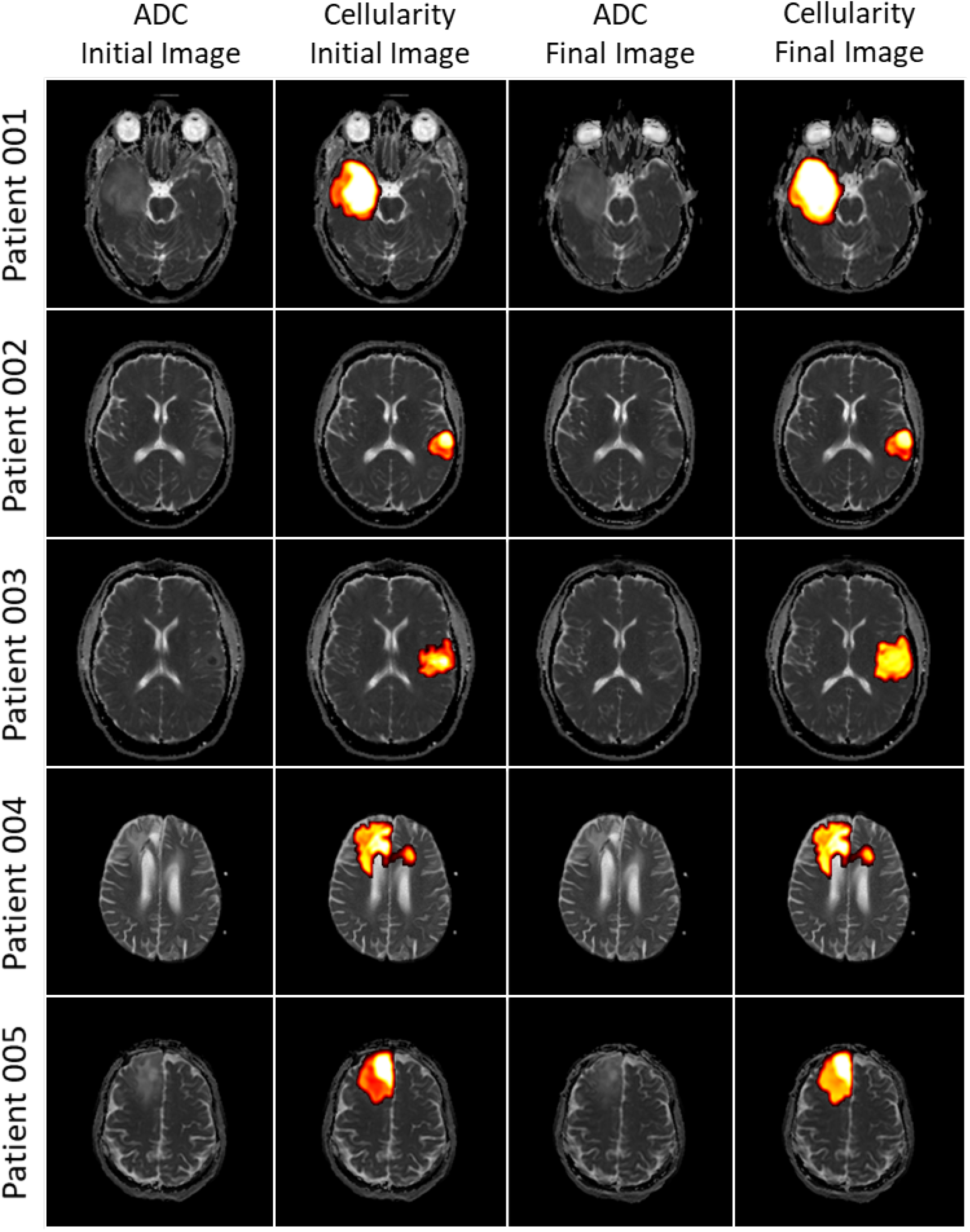
Representative cross sections of cellularity conversion from ADC and tumour segmentations for our patient dataset. Conversions were performed using equation (3). The first and third columns show the ADC at the initial and final image times respectively. The second and fourth columns show the ADC with the calculated cellularity superimposed at the initial and final image times respectively. Voxels that appear lighter (white and yellow) have a higher cell density and voxels that appear darker (black and red) have a lower cell density.

### 2.5 Deep Learning Parameter Estimation

Neural networks can be used for function inference; in other words, to copy the action of a target function on given inputs. If given *N* input-output pairs from a target function, 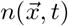, a neural network can be trained to approximate the function by minimizing the mean squared error between the expected and predicted value of that function

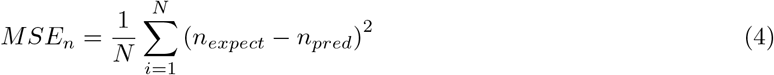

where *n*_*expect*_ is the true value of the function and *n*_*pred*_ is the value of the function predicted by the neural network. While this works well in theory and for sufficiently simple functions, in practice, it tends to be less successful than hoped. This is because in most real-world examples (and especially for examples in medicine), functions tend to be complicated and data tends to be scarce and noisy - which makes training a function inference network challenging or impossible. However, in such situations, there is often more known information than just the raw data points, though incorporating this existing system knowledge into a neural network is not a straightforward task. In the problem examined in this work, we can supplement the data contained in the MRIs with the knowledge contained in the PI model.

Our method for incorporating the PDE into the neural network optimization is motivated by the concept of physics-informed neural networks (PINNs) developed in a series of papers and a patent by Raissi et al [29, 30, 31]. The main idea is to create a neural network which infers the cell density function 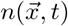 and train that network by using a loss function which incorporates a quantification of the satisfaction of the PI model by the predictions of the inference neural network. Through several examples in the original papers explaining PINNs, it is shown that this formulation results in the network converging more accurately and more efficiently than by using the data alone. This is not surprising, since by altering the loss function in this way, more information has been added from which the neural network can learn. Of particular importance, it is also shown that the optimization remains remarkably robust when noise is incorporated into the known data.

To do this, the PI model is discretized using a Runge-Kutta time stepping algorithm with *q* stages (where *q* is chosen to be the number of intermediate points at which the solution is predicted). Then, the Runge-Kutta scheme is rewritten with the initial and final solutions as functions of the *q* stages,

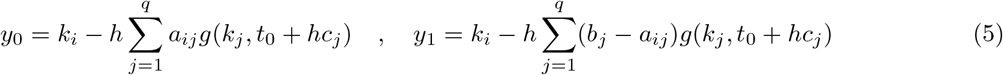

where *k*_*i*_ is the *i* th Runge-Kutta stage (*i* ∈ [0, *q*]), *a*_*ij*_, *b*_*i*_, and *c*_*i*_ are the Runge-Kutta scheme parameters, *h* is the time between the initial and final times, and *g* is the right hand side of the PI model (equation (1)) rewritten as 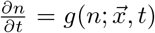. With this formulation, the network is set up to receive 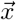 as input and output the Runge-Kutta stages *k*_1_, *k*_2_, …, *k*_*q*_. Evaluating the prediction of the initial and final data snapshots using these stages and the Runge-Kutta scheme, the loss function can be written as:

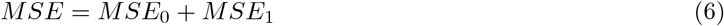

where

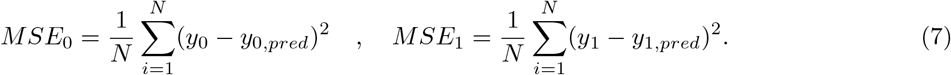

In the above, *N* is the number of data points at each of the two times, *y*_0_ and *y*_1_ are the known solutions at the initial and final times, and *y*_0,*pred*_ and *y*_1,*pred*_ are the solutions at the initial and final times predicted by the neural network. As the network is trained, MSE approaches zero and the network weights converge to the values which best mimic the action of the cell density function.

The key capability of this style of network optimization, however, is that parameters appearing in the PDE can be estimated during the training of the network. In the same way that the network optimizes the value of its weights, it can iteratively optimize the PDE parameters to find the values which best fit the data. This allows the network optimization to act as not only a method for solving the PI model, but also as a parameter estimation technique. Furthermore, this parameter estimation technique incorporates the entirety of the 3D tumour cellularity at both imaging times into its predictions.

## 3 Results

### 3.1 Synthetic Tumours

Before applying our deep learning model to the processed patient data, we test and showcase its capabilities by applying it to synthetic tumours. Synthetic tumours are computationally generated tumour cellularity profiles for which the full cellularity progressions and PI model parameter values are known. By first applying our deep learning model to these tumours, we can evaluate the model’s accuracy.

To generate a synthetic tumour, an initial condition is first selected. In the example shown in this section, we choose a Gaussian initial cellularity given by

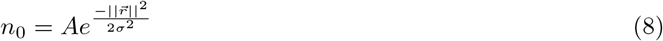

where we choose *A = n*_*max*_ = 1 and the standard deviation is chosen to be σ = 2mm. More complicated initial conditions can easily be chosen, but here we choose a simple distribution to clearly exhibit the model capabilities. We then select known values for the PI model parameters, *D* = 0.001mm^2^/day and *r* = 0.03/day, for this example. We then computationally solve the PI model using the chosen initial condition and model parameters over a time period of 50 days to arrive at a new cellularity profile for the tumour. We solve the PI model PDE using a finite element method implemented with the FEniCS project finite element solving software [2, 15].

The deep learning model accepts the initial and final cellularity profiles as input and outputs estimates for *D* and *r* as well as an estimate of the full cellularity progression between the two times known times. Since the true parameter values and growth curve are known, we can compare the predictions of the deep learning model to the known data to evaluate the model accuracy. This comparison can be seen in figure 4. We choose to predict the intermediate cellularity solution at 500 time points, meaning that we set up our numerical scheme to contain *q* = 500 Runge-Kutta stages. In the deep learning optimization, we utilize an Adam optimization over 30000 iterations and assume a constant learning rate of *η* = 0.001. For this example, the initial and final cellularity profiles were 3D arrays of size (31, 31, 31). Observe that the predicted cellularity matches the exact cellularity with high accuracy, and that the estimated values for *D* and *r* well approximate the true values: the relative error for *D* is 3.4% and for r is 1.6%.

**Figure 4:**
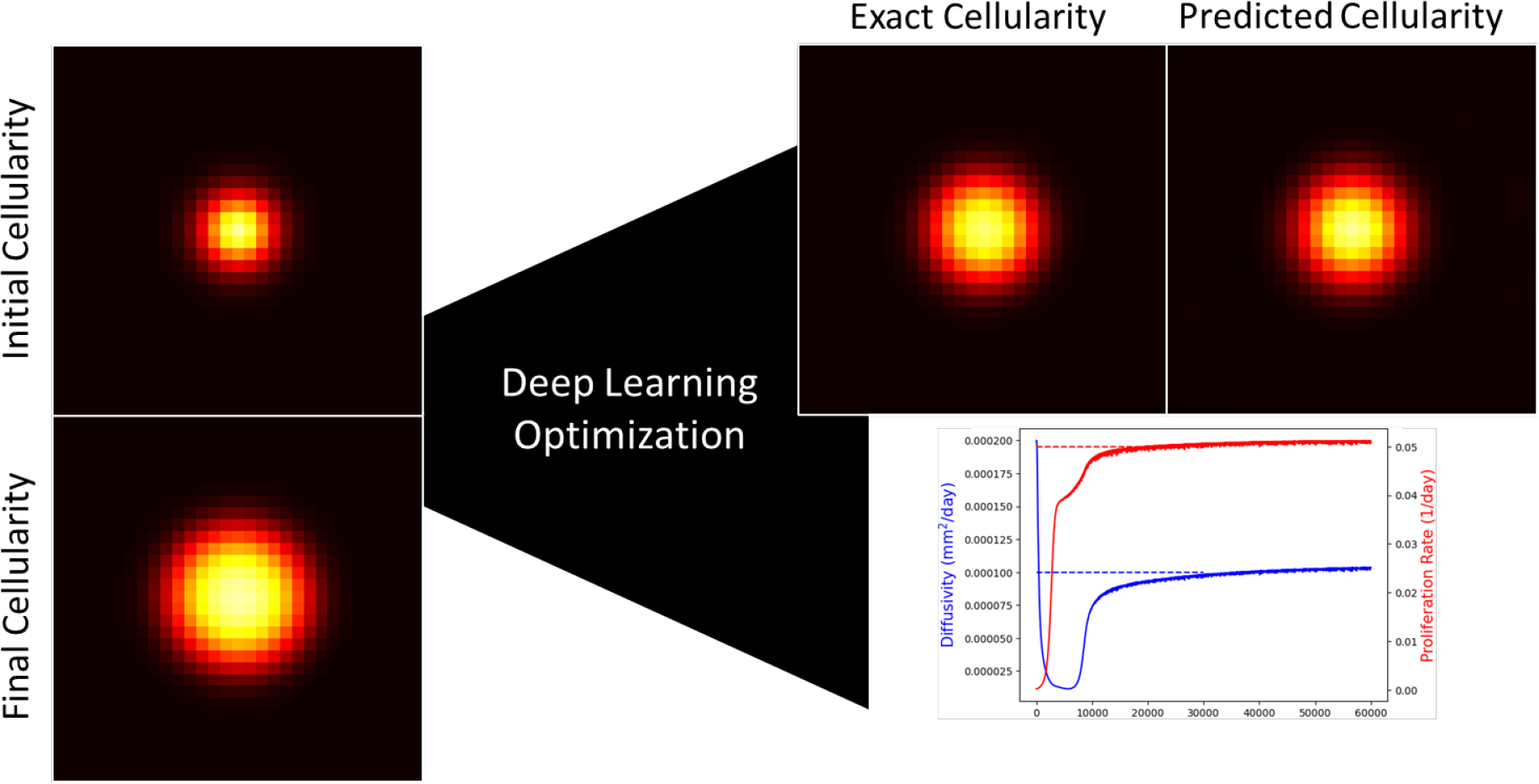
Results for the deep learning optimization of the synthetic tumour case. On the left, the initial and final cellularity profiles given as input to the deep learning model are shown. On the top right, a comparison of the exact and predicted cellularity at an intermediate time, halfway between the initial and final images is shown, showcasing the accuracy of the model in predicting the cellularity distribution. On the bottom right is the convergence of the predicted parameter values throughout network training. The flat dotted lines are the known, chosen values of the parameters (*D* = 0.0001mm^2^/day and *r* = 0.05/day).

### 3.2 Application to Patient Data

With confidence in the deep learning model after testing it on synthetic tumours, the model can be applied to the patient dataset. For each patient’s pair of cellularity profiles, we apply the deep learning model to estimate the values of the PI model parameters *D* and *r*. As the underlying data is not generated exactly as it was in the synthetic case, it takes far more iterations in order for the model to converge. Each patient is optimized over either 1 million or 1.5 million Adam iterations. The learning rate is chosen to start at *η* = 0.01 and decay by 2% every 5000 iterations.

See the resulting parameter estimates in table 2 and the predictions for the cellularity at intermediate times in figure 5. From estimated proliferation rate values, approximations for the doubling time of the tumour cell number can be calculated. Of course, the PI model assumes a logistic growth law, so these doubling time estimates are not true doubling times as they would be for an exponential growth law. Nevertheless, observe that the range of predicted doubling times is large. For example, patient 003 has a predicted doubling time of ≈ 10 days whereas patient 004 has a doubling time of over 600 days. Furthermore, notice how these estimates correspond with the imaging and cellularity profiles in figure 3. For example, the growth during the 51 days between images for patient 003 is significant, leading the model to predict a high proliferation rate. In contrast, the difference between the cellularity profiles for patient 004 appears small and the time between these images is 111 days, leading the model to converge to a small value of r and therefore large doubling time.

**Table 2:** Estimates for the PI model parameters and approximate tumour cell number doubling times for each of the patient data cases as predicted by our deep learning model.

| Patient ID | Estimated $D$ (mm<sup>2</sup>/day) | Estimated $r$ (1/day) | Approx. Doubling Time (Days) |
| --- | --- | --- | --- |
| 001 | 0.00094 | 0.02267 | 30.58 |
| 002 | 0.02538 | 0.00663 | 104.55 |
| 003 | 0.01173 | 0.06828 | 10.15 |
| 004 | 0.00000 | 0.00111 | 624.46 |
| 005 | 0.00099 | 0.00637 | 108.81 |

**Figure 5:**
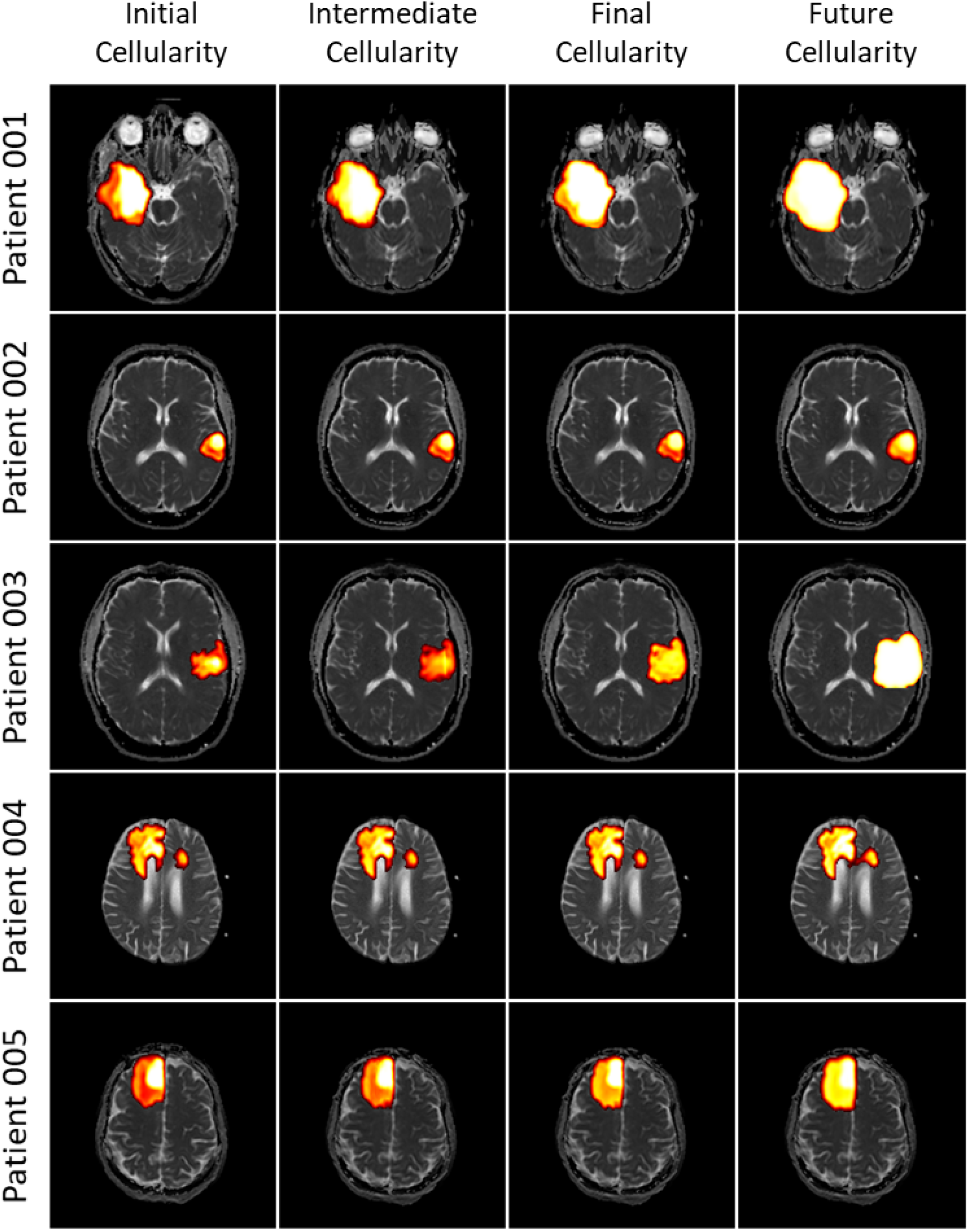
The results of the deep learning optimization model applied to the five patients in our dataset. The cross sections shown are representative slices of the 3D tumours. The cellularity is superimposed on the ADC at four times (from left to right): the initial imaging time, halfway between the two imaging times, the final imaging time, and 90 days after the final imaging time. The initial and final time images are from the data, while the intermediate time and future time cellularities are calculated by the model.

After obtaining the patient-specific estimates for the PI model parameters, they can be used to predict future tumour progression. To do this, we take the final cellularity profile for each patient and simulate the solution of the PI model with these parameter values. Like for the synthetic tumour case, solving the PI model is done using the FEniCS project solver [2, 15]. The results of this tumour progression projection can also be seen in figure 5.

## 4 Conclusion

In this work, we develop a pipeline that utilizes a deep learning model to make patient-specific estimates of the parameters of the well-known PI model for brain tumour progression and use these estimates to make personalized growth projections. The pipeline relies on knowledge of five MRI sequences, including T1, T1-GAD, T2, T2-FLAIR, and DWI - the first four of which are used in image segmentation and the final of which is used to derive a map of tumour cellularity. This method allows for more accurate and personalized predictions of tumour growth and potential treatment response than were otherwise possible, which clearly has theoretical and clinical utility. A particular advantage of this method is that it requires data only from the patient about whom predictions are being made. This sidesteps the major hurdle of requiring a large existing dataset which commonly plagues the application of machine learning models to problems in medicine.

While our method was capable of deriving reasonable predictions, there are several parts of our work that could be improved or generalized. First, no segmentation algorithm is perfect, and improvements to segmentation accuracy would directly lead to more accurate deep learning predictions. Luckily, including different segmentation algorithms into our pipeline is a simple process - meaning that as benchmark accuracy of brain tumour segmentation algorithms continues to improve, these gains can be seamlessly incorporated into our optimization. Our choice of FeTS as the segmentation algorithm has the advantage of classifying tissue into one of several categories; though since brain tumour segmentation is not the focus of this paper, we do not investigate and compare other potential choices. Second, clearly more research is necessary into how best to derive tumour cellularity from imaging. Since our conversion method is able to utilize the voxel-by-voxel tissue classification, we believe that the resulting cellularity profiles are likely more realistic than the standard method. However, more in depth studies in the literature leading to more accurate rules for cellularity derivation would be of tremendous use to improving the accuracy of our work. Furthermore, changes in this regard could similarly be seamlessly incorporated. The most significant source of error in our model however likely is due to differences in the MRI machine calibration and operator protocols. As has been noted elsewhere, standardization across image acquisition protocols is clearly necessary in order to advance AI in medicine [6, 7, 9, 14, 24]. When considering any of the above potential sources of error however, it is important to recall that our deep learning optimization is very robust with respect to noise within the data (as is explored in depth in [30, 29, 31]). Given this, even with changes to any of these parts of our procedure, changes to the parameter estimates are likely to be small. This lack of sensitivity to data noise is a key advantage of our method.

Though the PI model is a well-established tool for modelling the progression of brain tumours, other more sophisticated models could provide benefits. Our deep learning model could also easily be applied to a different model equation (or equations) to estimate the parameters appearing in that model. For example, the PI model could be augmented to include the effect of treatment by adding terms to the right hand side of equation (1) as has been done in many other works [19,20, 28,32, 33]. This would allow for the analysis of patients who have treatment between their images as well as the potential to make patientspecific estimates of mathematical parameters related to treatment such as the radiobiological parameters in the linear-quadratic model. Additionally, works using the PI model [25, 26, 40] often consider a spatially-dependent diffusivity,

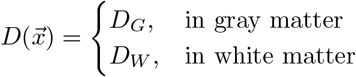

where differences between gray and white matter are considered. This could also be incorporated into our model by obtaining a map classifying brain voxels into gray and white matter, then adding a third parameter for the deep learning model to optimize. There is also the possibility of pretraining the deep learning model on a larger set of patients prior to patient-specific training. This could reduce the computational expense and potentially lead to more accurate predictions. Finally, recent works [18, 44] have noted differences in gliomas based on patient sex and have called for the inclusion of this information in analysis. For our work, it would be interesting to observe the differences in parameter predictions for male and female patients, though this would require a much larger dataset than we have used in order to be meaningful. This will be considered in future studies.

We hope that this work serves as an example of the benefits of deep learning applied to the analysis of medical imaging. Techniques such as this can act in addition to the standard clinical workflow, providing clinicians with evidence-based projections which incorporate patient data that can be used to aid in disease management.

## Acknowledgements

The financial support from the Natural Sciences and Engineering Research Council of Canada (NSERC) (CM and MK) is gratefully acknowledged.

## Notes

### Competing Interest Statement

The authors have declared no competing interest.

